# Genomic analysis reveals a polygenic architecture of antler morphology in wild red deer (*Cervus elaphus*)

**DOI:** 10.1101/2021.04.16.440189

**Authors:** Lucy Peters, Jisca Huisman, Loeske E.B. Kruuk, Josephine M. Pemberton, Susan E. Johnston

**Affiliations:** Institute of Evolutionary Biology, School of Biological Sciences, University of Edinburgh, Edinburgh, EH9 3FL, United Kingdom; Research School of Biology, The Australian National University, Canberra, Australia

## Abstract

Sexually-selected traits show large variation and rapid evolution across the animal kingdom, yet genetic variation often persists within populations despite apparent directional selection. A key step in solving this long-standing paradox is to determine the genetic architecture of sexually-selected traits to understand evolutionary drivers and constraints at the genomic level. Antlers are a form of sexual weaponry in male red deer. On the island of Rum, Scotland, males with larger antlers have increased breeding success, yet there has been no response to selection observed at the genetic level. To better understand the underlying mechanisms of this observation, we investigate the genetic architecture of ten antler traits and their principle components using genomic data from >38,000 SNPs. We estimate the heritabilities and genetic correlations of the antler traits using a genomic relatedness approach. We then use genome-wide association and haplotype-based regional heritability to identify regions of the genome underlying antler morphology, and an Empirical Bayes approach to estimate the underlying distributions of allele effect sizes. We show that antler morphology is heritable with a polygenic architecture, highly repeatable over an individual’s lifetime, and that almost all aspects are positively genetically correlated with some loci identified as having pleiotropic effects. Our findings suggest that a large mutational target and pleiotropy with traits sharing similar complex polygenic architectures are likely to contribute to the maintenance of genetic variation in antler morphology in this population.

## Introduction

Sexually-selected traits show a great variety and complexity across the animal kingdom, ranging from traits and behaviours that increase attractiveness, to those which increase intra-sexual competitiveness for access to mates (Andersson, 1994). Sexual traits are typically under strong selection (Kingsolver *et al.*, 2001), with phenotypic differences between related species suggesting that they can evolve rapidly, with downstream consequences for other phenomena such as adaptation, speciation and extinction probability (Lorch *et al.*, 2003; Ritchie, 2007; Servedio & Bürger, 2014; Wilkinson *et al.*, 2015; Martínez-Ruiz & Knell, 2017). Theory predicts that such strong sexual selection will reduce genetic variation within populations, yet empirical studies often show that sexual traits have substantial underlying genetic variation despite evidence of selection (Pomiankowski & Moller, 1995; Kotiaho *et al.*, 2001; Kruuk *et al.*, 2002; Svensson & Gosden, 2007). This contradiction presents an evolutionary paradox, for which several explanations have been proposed. These include differences in selection between the sexes, developmental stages or environmental conditions (Bourret *et al.*, 2017; Barson *et al.*, 2015), phenotypic plasticity (Charmantier & Gienapp, 2014), condition dependence (Dugand *et al.*, 2019) and trade-offs with survival (Johnston *et al.*, 2013). In addition, these observations could be due to genetic correlations with traits under opposing selection gradients and linkage disequilibrium between causal loci and deleterious alleles (Lande, 1982; Lande & Arnold, 1983; Connallon & Hall, 2018). Quantitative genetic studies have provided some insight into these different explanations, through estimating the relative contributions of additive genetic (i.e. the heritability, *h*^2^) and environmental effects to phenotypic variance, as well as examining phenotypic and genetic correlations with other traits, including fitness (Emlen, 1994; Griffith *et al.*, 1999; Kruuk *et al.*, 2002; Robinson et *al.,* 2006). However, a key limitation of most studies to date is that the genetic architecture of sexual traits is generally unknown - that is, the underlying loci, their number, distribution and relative effect sizes, and the extent of pleiotropy, epistatis and other interactions (Timpson et *al.,* 2018; Chenoweth & McGuigan, 2010). By identifying the genetic architecture of sexual traits, we can better understand the underlying molecular mechanisms and evolutionary processes that drive their variation (Dobzhansky, 1971; Lewontin *et al.*, 1974; Kuijper *et al.*, 2012; Wilkinson *et al.*, 2015).

Recent genomic advances in natural populations have led to a number of studies characterising genetic architectures using genome-wide association studies (GWAS, reviewed in Santure & Garant 2018). Yet, relatively few studies exist for sexually selected traits, with much of the focus on discrete traits with Mendelian or relatively simple genetic architectures (Johnston *et al.*, 2011; Barson *et al.*, 2015; Hendrickx *et al.*, 2021). In these rare cases, mapping specific genomic variants associated with sexual trait variation can allow investigation of sex, age and environment-specific effects at individual loci. As such, they have revealed compelling cases of heterozygote advantage due to trade-offs between reproductive success and survival (Johnston *et al.*, 2013), or due to differences in optimal trait expression between the sexes (Barson *et al.*, 2015). However in most cases, sexual traits are likely to have oligogenic or polygenic architectures (i.e. moderate to large numbers of underlying loci), particularly in cases where they are condition dependent (Rowe & Houle, 1996). One issue is that as the number of loci increase and their relative effect sizes decrease, it becomes more difficult to implicate individual loci in trait variation; for example, in heights of people of European ancestry, only a fraction of the loci underpinning variation has been identified (Yengo *et al.*, 2018). On the other hand, being able to determine that a trait has a polygenic architecture can still shed light on how sexual traits evolve for the following reasons. First, polygenic traits present a large mutational target, contributing to the maintenance of genetic variance via the introduction of new variants (Rowe & Houle, 1996). Second, selection induced allele frequency changes at a great number of loci is expected to result in a rapid change in trait mean, and thus trait evolution, which is sustained by the aforementioned large mutational input, leaving the distribution of genetic effects unperturbed (Barton *et al.*, 2017; Sella & Barton, 2019). Third, pleiotropy and/or linkage between loci could maintain variation through conflicts between traits sharing a similar or linked polygenic architecture (Ruzicka *et al.*, 2019). Therefore, studies of sexual traits should aim to identify specific genetic variants with large effects on phenotype, and should also aim also to determine the distribution of polygenic effect sizes and the degree to which these underlying loci are shared between traits. This will not only shed light on potential evolutionary processes and mechanisms in empirical studies, but will also inform the mechanistic details of theoretical models to allow better assumptions to account for the complexities of natural populations (McNamara & Houston, 2009; Wilkinson *et al.*, 2015).

Antlers are a form of sexual weaponry in deer (Cervidae) that are generally only present in males and are shed and regrown annually (Davis *et al.*, 2011). They are used as weaponry in intra-male competition for access to females, with larger antlers often associated with increased reproductive success (Kruuk *et al.*, 2002; Malo *et al.*, 2005), yet antler weights and dimensions are often moderately heritable in both wild and captive populations (Lukefahr & Jacobson, 1998; Williams *et al.*, 1994; Wang *et al.*, 1999; Van Den Berg & Garrick, 1997; Kruuk *et al.*, 2002, 2014; Jamieson *et al.*, 2020). In male red deer (*Cervus elaphus*) on the island of Rum, Scotland, there is directional selection for increased antler weight and number of points (known as “form”), yet both are substantially heritable (*h*^2^ = 0.38 & 0.24, respectively) with no phenotypic response to selection observed over a 30 year study period (Kruuk *et al.*, 2002, 2014). Whilst both antler weight and form are positively genetically correlated, the selection gradients on the genetic components were estimated as zero and negative, respectively (Kruuk *et al.*, 2002, 2014). This suggested that selection for antler weight is environmentally driven, whereas negative selection on antler form is constrained by genetic associations with a trait that is genetically unresponsive to selection, meaning that genetic constraints may contribute to the maintenance of genetic variance. Alternatively, the association with breeding success may be driven by indirect correlations with environmental variables (Kruuk *et al.*, 2014). To better understand the mechanisms constraining selection, a logical next step is to determine the genetic architecture of antler morphology. As antlers present a multi-dimensional phenotype, adding more information from different measures may contribute to understanding potential evolutionary conflict and constraints within the antler and to characterise the specific genetic variants underpinning heritable variation (Chenoweth & McGuigan, 2010).

In this study, we used an extensive antler morphology data set with 948 to 3972 observations in 336 to 891 unique males and genomic data from 38,000 polymorphic SNPs to estimate the heritability and genetic correlations of ten antler traits using genomic relatedness matrices. We then use genome-wide association and haplotype-based regional heritability to identify regions of the genome underlying antler morphology, and an Empirical Bayes approach to estimate the underlying distributions of allele effect sizes. We show that antler morphology is heritable with a polygenic architecture, and that almost all aspects are positively genetically correlated with some loci identified as having pleiotropic effects. Our findings suggest that genetic variation in antler morphology is maintained via a large mutational target and pleiotropy with traits sharing similar complex polygenic architectures in the red deer population.

## Methods

### Study system

The red deer study population is situated in the North Block of the Isle of Rum, Scotland (57°02’N, 6°20’W) and has been subject to individual monitoring since 1971 (Clutton-Brock *et al.*, 1982). Deer calves are marked with ear tags shortly after birth to enable recording of detailed life histories of individuals. DNA is routinely extracted from neonatal ear punches, post-mortem tissue and/or cast antlers (see Huisman *et al.* 2016). A pedigree of 4,515 individuals is available for the population, and was previously constructed using single nucleotide polymorphism (SNP) data in the R package Sequoia (Huisman, 2017). Research was conducted following approval of the University of Edinburgh’s Animal Welfare and Ethical Review Body and under appropriate UK Home Office licenses.

### Antler measures

Male red deer cast and regrow their antlers every year from the age of one or two (Kruuk *et al.*, 2002). Ten antler measures are routinely taken from cast antlers and antlers from deceased individuals between 1971 and 2017 (see Figure 1 and Table 1 for full details and sample sizes of each measure). All measures of length were taken following the curve of the antler, and measures of circumference were taken at the narrowest point between antler tines (points). Total antler length was defined as the distance from the coronet (base) to the furthest point of the antler. All length and circumference measures were taken in centimetres. Antler weight was measured as the total dry mass of the antler in grams. Antler form was defined as the number of tines, and as this trait can be determined visually on living deer, was determined from observations in the field and therefore has the greatest sample size (3972 observations from 891 stags). Where measurements from both the left and right antlers were available, the mean was taken. Individual measurements were excluded if the antler part was broken and antler weight was discarded if any part of the antler was broken. Only antlers from stags aged 3 years or older were considered, as cast antlers recovered in the field can be reliably assigned to known individuals by their shape from this age onwards (Kruuk *et al.*, 2002, 2014).

**Figure 1:**
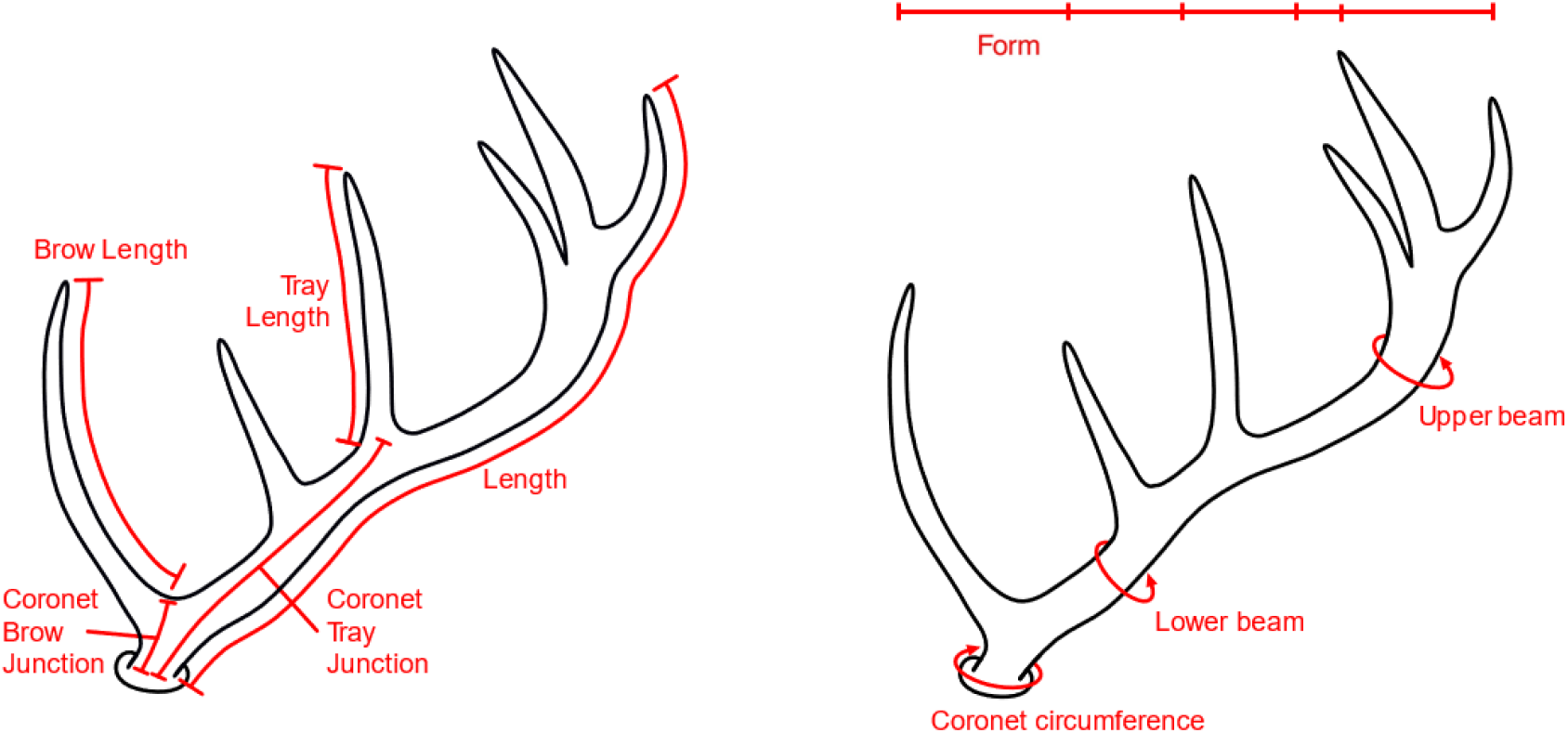
A schematic of the antler measures used in this study (measured in cm). Antler weight was also measured in g. Further details for each measurement are given in the main text and in Table 1.

**Table 1:** Summary statistics of all 10 antler measures as shown in Figure 1 in SNP genotyped males. *N Obs* is the number of observations, *N IDs* is the number of unique individuals for each measure. All lengths and circumferences were measured in *cm,* weight was measured in *g. SD* denotes the standard deviation.

| Trait | N Obs | N IDs | Mean | SD |
| --- | --- | --- | --- | --- |
| Antler Length | 1060 | 444 | 65.168 | 12.029 |
| Coronet Circumference | 1224 | 471 | 15.626 | 2.013 |
| Lower Beam Circumference | 1204 | 460 | 10.504 | 1.533 |
| Upper Beam Circumference | 1123 | 436 | 9.446 | 1.358 |
| Coronet-Brow Junction | 1162 | 466 | 5.720 | 1.025 |
| Coronet-Tray Junction | 1184 | 455 | 27.313 | 5.398 |
| Brow Length | 1139 | 457 | 20.568 | 5.461 |
| Tray Length | 948 | 388 | 14.484 | 4.177 |
| Antler Weight | 1003 | 336 | 649.858 | 243.098 |
| Form (Total Number of Points) | 3972 | 891 | 4.498 | 1.072 |

### Genomic data-set

DNA samples from 2,870 individuals have been genotyped at 51,248 SNP markers (Huisman *et al.*, 2016) on the Cervine Illumina BeadChip (Brauning *et al.*, 2015) using an Illumina genotyping platform and Illumina GenomeStudio software (Illumina Inc., San Diego, CA, USA). All SNPs on the Cervine Illumina BeadChip are named based on their synteny with the cattle genome BTA vUMD 3.0 (e.g. SNP ID *cela1_red_15_1479373* is orthologous to position 1479373 on cattle chromosome 15). In addition, a linkage map specific to the Rum population is available, with 38,083 SNPs assigned to linkage groups corresponding to the 33 deer autosomes and X chromosome (Johnston *et al.*, 2017). Quality control was carried out in PLINK v1.9 (Chang *et al.*, 2015) with the following thresholds: SNP genotyping success >0.99, minor allele frequency >0.01 and individual genotyping success >0.99. Further quality control of mapped SNPs and X-linked markers (i.e heterozygous state in males) was conducted using the *check.marker* function with default thresholds in the R library GenABEL v1.80 in R v3.4.2 (Aulchenko *et al.*, 2007). The final SNP dataset consisted of 2,138 individuals and 38,006 markers. Genome-wide linkage disequilibrium (LD) was calculated between all SNPs within 1Mb of each other using Spearman’s Rank correlation (*ρ*). Based on a linear regression of *ρ* on the log base pair distance between SNPs, LD decayed at a relatively low rate of 0.031 *ρ* per Mb (SE = 5.56×10 ‘, Figure S1).

### Principal components of antler measures

A principal component analysis (PCA) was conducted to create a second dataset combining information from the different antler measures, while also increasing the differentiation among the different principal components (PCs). As PCA does not allow for missing data, we imputed missing antler measures using the Bayesian *bpca* algorithm in the packages pcaMethods v1.7 in R v3.4.2 (Stacklies & Redestig, 2018; Oba *et al.*, 2003). We used the default settings which assumes a flat prior distribution for imputation, and the most appropriate number of PCs was determined using the *kEstimate* function. To improve imputation accuracy, the form was split into the number of tines on the lower beam and upper beam, respectively, resulting in 11 antler measures. Imputation accuracy was quantified by calculating the error of prediction (*E*) from a complete subset of the data with no missing values and the same subset with randomly missing data at a similar level to the whole data set (~ 9%). The error of prediction was calculated as follows:

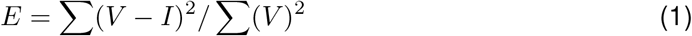

where *I* refers to the imputed values of the data subset with missing values and *V* to their counterpart in the complete data subset (Stacklies & Redestig, 2018). Our analysis fount that the imputation of missing values using a Bayesian PCA approach achieved high accuracy (*E* = 0.015). To account for variation among antlers due to age structure, antler measure values were modelled using a linear model approach (following Pallares *et al.* 2014). All models had the same structure:

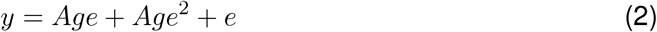

where *y* is a vector of the antler measure and *e* is a residual error term. Age effects were fitted as a fixed quadratic term to account for the non-linear change in antler measures with age (see also Nussey *et al.* 2009 and Kruuk *et al.* 2002). All antler measures were modelled using a Gaussian distribution. All models were fitted using the *lm* function in R v3.4.2. The residuals of these models were then used in a standard PCA using the *prcomp* function in R and the scores of the PCs used as trait values in the downstream analysis.

### Estimating heritability using the animal model

All antler measures and PCs were modelled using a restricted maximum-likelihood (REML) approach within the mixed ‘animal model’ framework (Henderson, 1975) in ASREML-R v4.0 (Butler *et al.*, 2009) in R v3.4.2. The animal model estimates the effect sizes of fixed effects and partitions phenotypic variance (*V_P_*) into several random effects, including the variance attributed to additive genetic effects (*V_A_*). Previous studies have estimated *V_A_* using a pedigree relatedness matrix (Kruuk *et al.*, 2002, 2014); here, we wished to compare estimates of *V_A_* using both pedigree and genomic relatedness information. Therefore, all models were carried out estimating *V_A_* in one of two ways: 1) using a numerator relationship matrix *A* based on the pedigree, using the *ainv* function in ASReml-R; and 2) using a genomic relatedness matrix (GRM) calculated using autosomal SNPs (N = 37,271) in the *--make-grm function* in GCTA v1.24.3 (Yang *et al.*, 2011a). The GRM was adjusted to assume similar frequency spectra of genotyped and causal loci with the argument *--grm-adj 0.*

All 10 antler measures and 11 PCs were modelled in univariate animal models with the following structure:

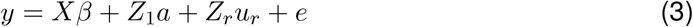

where *y* is a vector of the antler measure or PC, *X* is an incidence matrix relating individual measures to the vector of fixed effects *β*; *Z*_1_ and *Z_r_* are an incidence matrices relating individual measures to additive genetic and other random effects respectively; *a* is a vector of relatedness matrix *A* or GRM; *u_r_* is a vector of additional random effects; and *e* is a vector of residual effects. Fixed effects included age in years as a both a linear and quadratic term for the antler measures and an intercept only for the PC models (as age structure was accounted for prior to PC estimation). Random effects included: the additive genetic effect; permanent environment (i.e. individual identity) to account for pseudoreplication due to repeated measures in the same individual; and birth year and year of antler growth to account for common environmental effects between individuals. The narrow sense heritability (*h*^2^) was calculated as *V_A_/V_P_*, where *V_P_* was defined as the sum of the variance attributed to all random effects, including the residual variance (Falconer & Mackay, 1996). The significance of fixed effects was calculated with a Wald test, while the significance of the random effects was tested using a likelihood ratio test (LRT) between models with and without random effect of interest (i.e. 2 × the difference between the model log-likelihoods, assuming a *χ^2^* distribution with 1 degree of freedom).

Bivariate models were run to determine genetic correlations between the 10 antler measures, with the following structure:

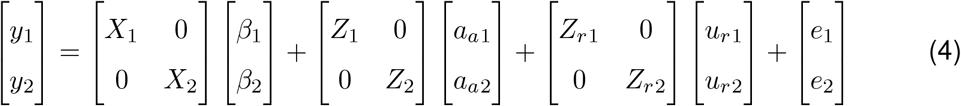

All variables are as defined in equation (3), with subscripts referring to antler traits 1 and 2, respectively. The GRM was used to model the additive genetic covariance. The genetic correlation *r*^2^ can be obtained from the genetic covariance, as *r_a_* = *cov_a_*(1,2)/*σ*_1_*σ*_2_, where *cov_a_*(1,2) stands for the covariance between trait 1 and 2, and *σ* represents the respective standard deviations for traits 1 and 2. The significance of *r_a_* was determined using an LRT as above, by comparing the model to another where *r_a_* was constrained to either zero or one.

### Genome-wide association studies

Genome-wide association studies (GWAS) were conducted in RepeatABEL v1.1 (Rönnegård *et al.*, 2016) implemented in R v3.4.2. First, the *prefitModel* function was used to fit a linear mixed model (without fixed SNP effects), specifying the same fixed and random effect structure as Equation 3. The resulting covariance matrix of the random effects was then input to the *rGLS* function, which fits each SNP genotype as an additive linear covariate. This approach accounts for population structure by fitting the GRM as a random effect and allows for repeated measures per individual. The significance of each SNP was determining using Wald tests, distributed as *χ*^2^ with 1 degree of freedom. These statistics were corrected for potential inflation due to population structure not captured by the GRM by dividing them by the genomic inflation factor *λ,* defined as the observed median *χ*^2^ statistic divided by the null expectation median *χ*^2^ statistic (Devlin & Roeder, 1999). This was done separately for each antler measure. After correction, the genome-wide significance threshold was set to a p-value of 1.42×10^-6^, equivalent to α= 0.05 after correcting for multiple-testing and accounting for non-independence due to LD between SNP markers (see Johnston *et al.* 2018).

### Regional heritability analysis

In addition to GWAS, we used a regional heritability approach to identify regions of the genome associated with antler trait variation. This method uses information from multiple loci to determine the proportion of *V_P_* explained by defined genomic regions, and has increased power to detect variants of small effect sizes and low minor allele frequencies (Yang *et al.*, 2011b; Nagamine *et al.*, 2012). Regions were defined using a ’sliding window’ approach with SNPs of known position on the Rum deer linkage map (Johnston *et al.*, 2017). Each window was 20 SNPs wide and overlapped the preceding window by 10 SNPs. If the last window in the linkage group contained less than 20 SNPs, the last 20 SNPs of that linkage group were taken instead. SNPs in linkage group 34, which corresponding to the X chromosome, were excluded from this analysis, as models using X-linked markers did not converge. This resulted in a total of 3,608 genomic windows. The contribution of each genomic region to *V_A_* and *V_P_* for each antler measure and PC was modelled as follows:

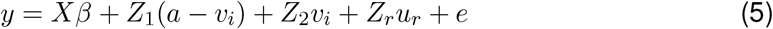

with variables defined as in Equation (3), but with the additive genetic components split into two terms: *Z*_1_(*a* – *v_i_*) and *Z*_2_*v_i_*, where *Z*_1_ is an incidence matrix of the GRM constructed based on all autosomal SNPs excluding those in window *i*, (*a* – *v_i_*) is the additive genetic effect excluding the window *i*; *Z*_2_ is an incidence matrix of the GRM constructed with only the SNPs in window *i* and *v* is the additive genetic effect of the window *i*. The significance of an association between a window *i* and an antler trait was determined using a LRT comparing models containing and omitting the term *Z*_2_*v_i_*. The distribution of *χ*^2^ statistics from the LRTs across all windows was corrected using the genomic control parameter *λ,* calculated using the same approach as above (Devlin & Roeder, 1999). A genome-wide significance threshold was calculated using a Bonferroni correction, where the *α* significance level (here 0.05) was divided by the number of effective tests. For this, we divided the number of windows by two to account for the overlap of half the number of total SNPs within each window, resulting in a significance threshold of P = 2.77×10^-5^.

### Estimation of SNP effect size distribution

We investigated the distribution of allele effect sizes and false discovery rates for all antler measures and PCs using the *ash* function in the R package ashR v2.2-32 (Stephens, 2016). This uses “adaptive shrinkage”, an Empirical Bayes method that uses the slopes and standard errors of the additive SNP effects from the GWAS models above to compute a posterior distribution of SNP effect sizes across all loci. This approach estimates the local false discovery rate (*lfdr*), which is the probability that the SNP effect is zero. The significance of a SNP effect was then determined by a local false sign rate (*lfsr*), defined as the probability of making an error when assigning a sign (positive or negative) to an effect, with a cut-off at *α* = 0.05. The prior distribution was specified to be any symmetric unimodal distribution when applying the *lfdr* estimation.

We then re-estimated the effect sizes of SNPs with the highest non-zero effects using adaptive shrinkage using the animal model framework (Equation 3) in ASREML-R v4.0 (Butler *et al.*, 2009). A maximum number of 10 SNPs per trait were taken. SNP genotype was fit as a two or three level factor and significance was tested using a Wald test. Fitting SNP genotype as a factor allowed the quantification of both the dominance deviation and the variance co-variance structure of the genotypes, which are needed to estimate the variance attributed to each SNP, calculated as follows (Falconer & Mackay, 1996):

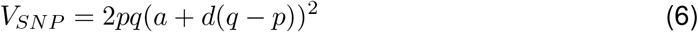

where *p* and *q* are the allele frequencies of alleles *A* and *B,* respectively; *a* is the additive genetic effect defined as the mid-point between the effect sizes of the genotypes *AA* and *BB*; and *d* is the dominance deviation defined as the difference between *a* and the effect size of the heterozygote *AB*. The proportion of *V_A_* attributed to a SNP was calculated as the ratio of *V_SNP_* to the sum of *V_SNP_* and the *V_A_* obtained from an animal model where the SNP effect was omitted. Standard errors of *V_SNP_* were estimated using the *deltamethod* function in the R library msm v1.6.7 (Jackson, 2011) in R v3.4.2.

### Data Accessibility Statement

Data for this study will be archived in a public repository upon manuscript acceptance. All results and data underlying the figures in this manuscript are provided as Supplementary Material. All scripts for the analysis are provided at https://github.com/Lucy-Peters/Red_deer_antler_genetic_architecture.

## Results

### Principal component analysis of antler measures

A principal component analysis (PCA) of 11 antler measures resulted in the maximum number of principal components (i.e. 11 PCs). The composition of the PCs showed that PC1, which explained around 41% of the variance, combined approximately equal amounts of information from all 11 measures, while PCs 2 to 11 explained increasingly less variance, mostly representing one or two antler measures (Figure 2).

**Figure 2:**
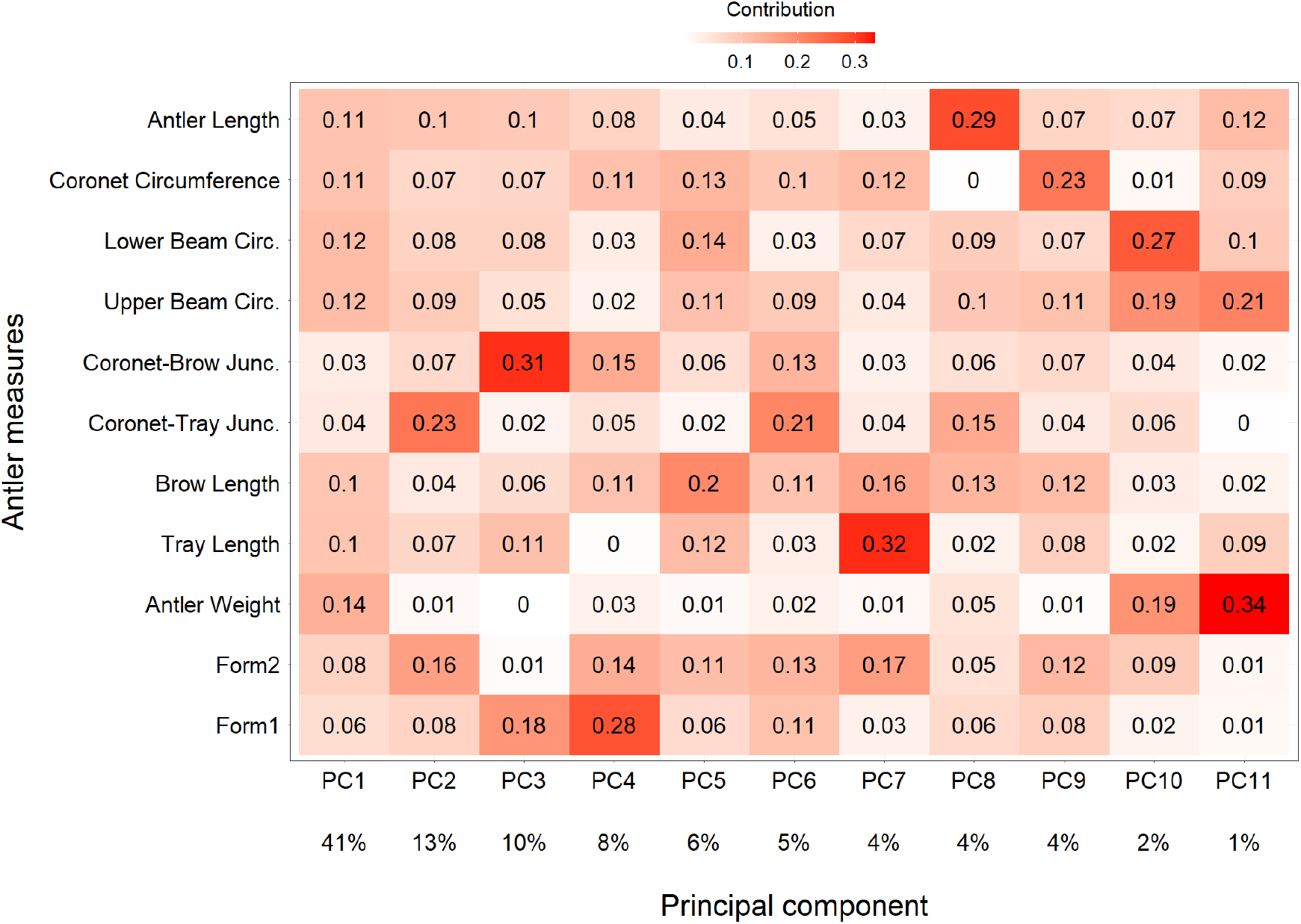
Heat-map showing the contribution (as a proportion) of each antler measurement to each of the 11 principal components.

### Animal models of antler measures

Antler measures were significantly heritable, with estimates ranging from *h*^2^ = 0.211 to 0.436 for the pedigree estimates, and *h*^2^ = 0.229 to 0.414 for the genomic estimates (Figure 3; Tables 2 & S1). Heritability estimates for antler weight, antler length, coronet circumference, brow length and form were generally consistent with previous findings by Kruuk *et al.* (2014, Figure 3). All antler PCs were also significantly heritable, although estimates decreased substantially for higher order PCs, which explained relatively small proportions of variance in antler morphology (Figure S2, Table S2). For all antler measures and PCs, confidence intervals between pedigree and genomic relatedness estimates were highly similar; there was no trend when comparing estimates of *h*^2^ for the same trait, suggesting that both the pedigree and genomic relatedness matrices capture the additive genetic variance to a similar degree in this population. Therefore, all results described from this point onwards are from models fitting a GRM, unless otherwise stated.

**Figure 3:**
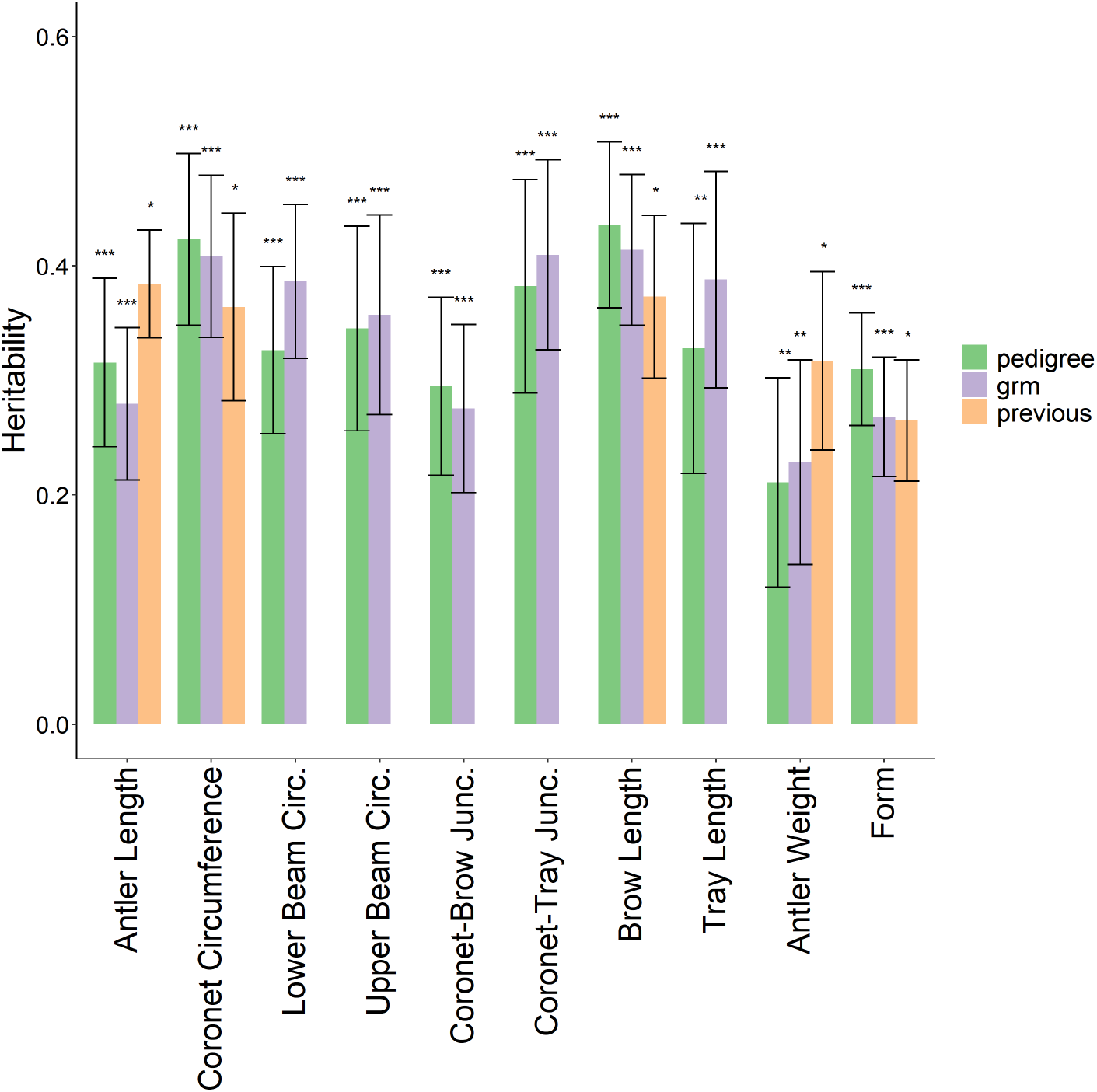
Heritability estimates for all 10 antler measures. Both estimates from the pedigree and the GRM models are shown as well as results from a previous study by Kruuk *et al.* (2014) for antler weight and form. \**P* ≤ 0.05, \*\**P* ≤ 0.01 and \*\*\**P* ≤ 0.001. Underlying data is provided in Table S1

**Table 2:** Proportions of phenotypic variance (*V_P_*) explained by random effects in animal models of the ten antler measures. The additive genetic effect was estimated using the GRM. Standard errors are given in brackets. Information on sample sizes and mean measures is provided in Table 1.

| Trait | $V_P$ | Additive Genetic | Permanent Environment | Birth Year | Growth Year | Residual | Repeatability |
| --- | --- | --- | --- | --- | --- | --- | --- |
| Antler Length | 61.155 | 0.279<br>(0.066) | 0.105<br>(0.058) | 1.04E-08<br>(0.014) | 0.222<br>(0.047) | 0.393<br>(0.035) | 0.385 |
| Coronet Circ. | 3.047 | 0.408<br>(0.071) | 0.128<br>(0.056) | 0.018<br>(0.018) | 0.243<br>(0.047) | 0.203<br>(0.020) | 0.554 |
| Lower Beam Circ. | 1.323 | 0.386<br>(0.067) | 0.096<br>(0.054) | 2.22E-07<br>(1.64E-08) | 0.211<br>(0.042) | 0.307<br>(0.027) | 0.483 |
| Upper Beam Circ. | 1.084 | 0.357<br>(0.087) | 0.280<br>(0.081) | 5.16E-08<br>(0.017) | 0.164<br>(0.035) | 0.199<br>(0.018) | 0.637 |
| Coronet-Brow Junc. | 0.872 | 0.275<br>(0.073) | 0.254<br>(0.068) | 0.026<br>(0.022) | 0.152<br>(0.037) | 0.292<br>(0.025) | 0.555 |
| Coronet-Tray Junc. | 20.930 | 0.410<br>(0.083) | 0.148<br>(0.072) | 0.001<br>(0.016) | 0.019<br>(0.009) | 0.422<br>(0.032) | 0.559 |
| Brow Length | 21.555 | 0.414<br>(0.066) | 0.068<br>(0.050) | 6.33E-09<br>(0.013) | 0.269<br>(0.050) | 0.250<br>(0.025) | 0.481 |
| Tray Length | 13.809 | 0.388<br>(0.094) | 0.273<br>(0.088) | 7.65E-08<br>(5.34E-09) | 0.028<br>(0.012) | 0.311<br>(0.028) | 0.661 |
| Antler Weight | 30727.710 | 0.229<br>(0.089) | 0.263<br>(0.086) | 0.023<br>(0.027) | 0.252<br>(0.050) | 0.233<br>(0.025) | 0.514 |
| Form | 0.907 | 0.268<br>(0.052) | 0.249<br>(0.046) | 0.024<br>(0.015) | 0.044<br>(0.011) | 0.416<br>(0.019) | 0.541 |

The permanent environmental effect (which includes dominance and epistatic effects) was generally significant for all antler measures, explaining up to 28.0% (upper beam circumference) of the phenotypic variance (Table 2). Year of antler growth explained a significant proportion of phenotypic variance for most antler measures and PCs (Tables 2 and S2, respectively). Conversely, birth year was not significant for any antler measure or PC; nevertheless, we retained this random effect in all models to account for potential cohort effects (Tables 2 & S2). Trait repeatabilities, calculated as the sum of contributions from the additive genetic, the permanent environment and birth year components, was high for all antler measures, ranging from 38.5% (antler length) to 66.1% (tray length; Table 2). Age as both linear and quadratic fixed effect terms was significantly associated with all antler measures (Table S3).

### Genetic correlations between antler measures

Most antler measures were positively genetically correlated, suggesting some degree of shared genetic architecture (Table 3). This was also reflected in the disproportionate contribution of all antler measures to PC1 (Figure 2). Antler weight was significantly positively correlated with almost all other measures (with the exception of coronet-tray junction), ranging from r^2^ = 0.40 for coronet-brow junction to r^2^ = 0.94 for upper beam circumference. The only significant negative genetic correlations were observed between coronet-tray junction and tray length, form and lower beam circumference (Table 3).

**Table 3:**
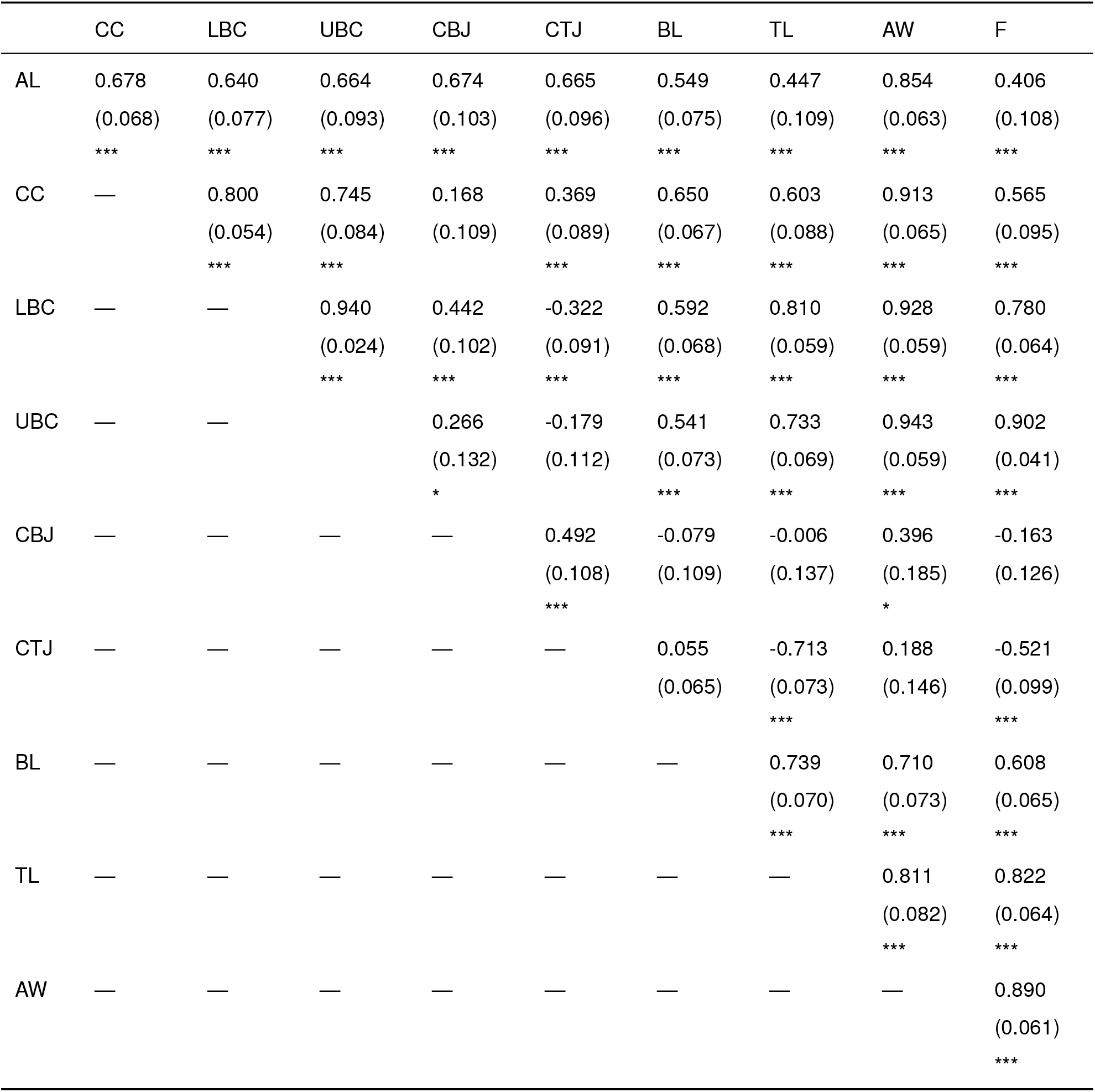
Genetic correlations among all 10 antler measurements. Standard errors are given in brackets, \**P* ≤ 0.05, \*\**P* ≤ 0.01 and \*\*\**P* ≤ 0.001. AL = Antler Length, CC = Coronet Circumference, LBC = Lower Beam Circumference, UBC = Upper Beam Circumference, CBJ = Coronet-Beam Junction, CTJ = Coronet-Tray Junction, BL = Brow Length, TL = Tray Length, AW = Antler Weight, F = Form.

### Genome-wide and regional heritability studies

No genomic regions were significantly associated with any antler measure or PC using GWAS (Figures 4 and S3, and Tables S4 and S5, respectively). The regional heritability analysis found no regions of the genome significantly associated with any antler measure or PC (Figures 5 and S4, and tables S6 and S7, respectively), with the exception of PC9, which was significantly associated with three overlapping windows (corresponding to a ~4.4 Mb region) on CEL linkage group 21. The most significantly associated window explained 16.8% (SE = 8.0%) of the phenotypic variance and 66.3% (SE = 22.0%) of the additive genetic variance. However, PC9 accounts for only 4% of overall phenotypic variance among all antler measures (see Figure 2). Homology with the cattle genome (version ARS-UCD1.2) suggested that there are a total of 19 coding regions within the region covered by all three significant windows, Details of SNPs within this region can be found in Table S8 and associated GO terms can be found in Table S9.

**Figure 4:**
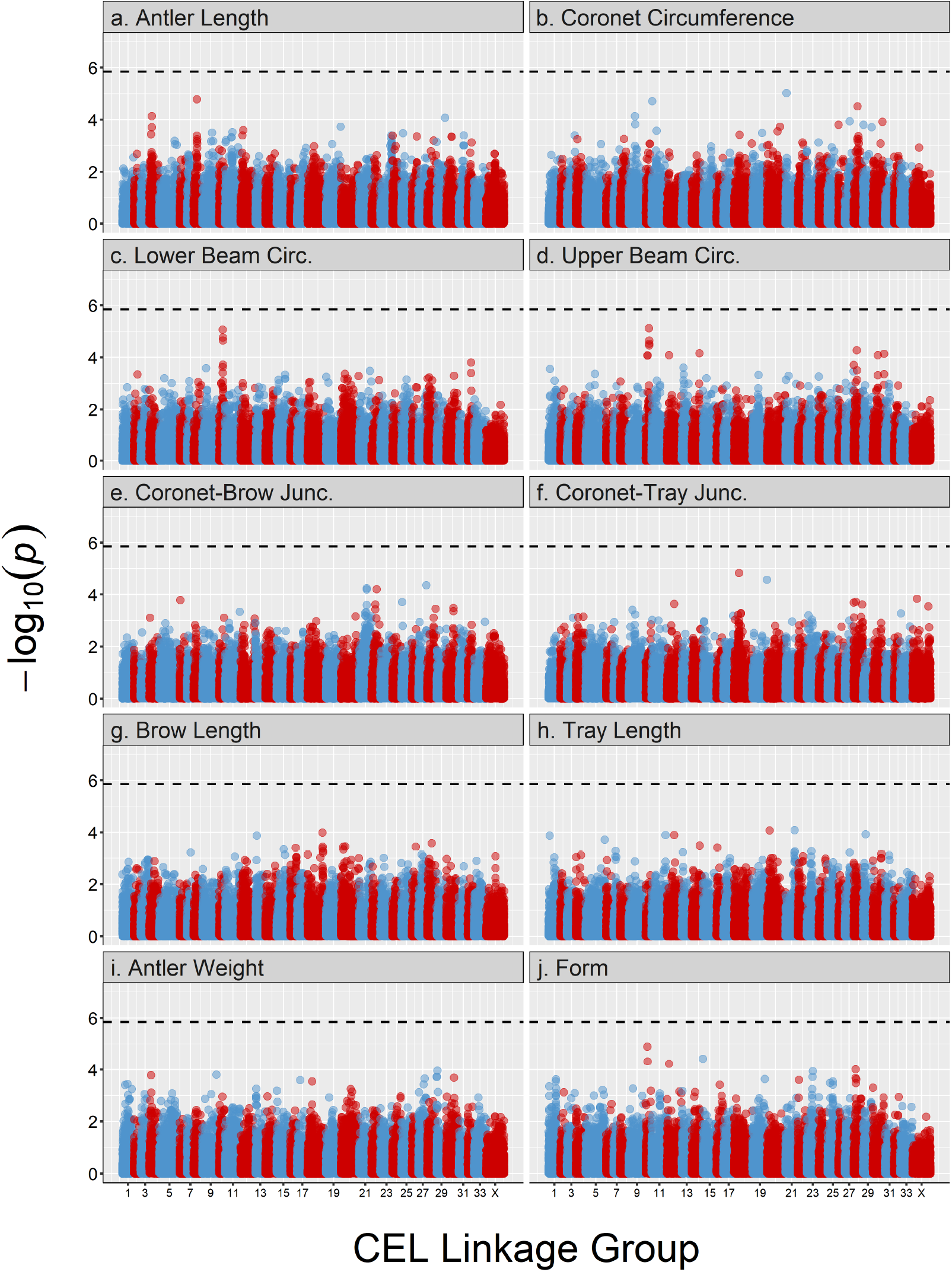
Genome-wide association study for antler measures. The dashed line indicates the significance threshold equivalent to *α* = 0.05. Points are colour-coded by chromosome. Underlying data is provided in Table S4.

**Figure 5:**
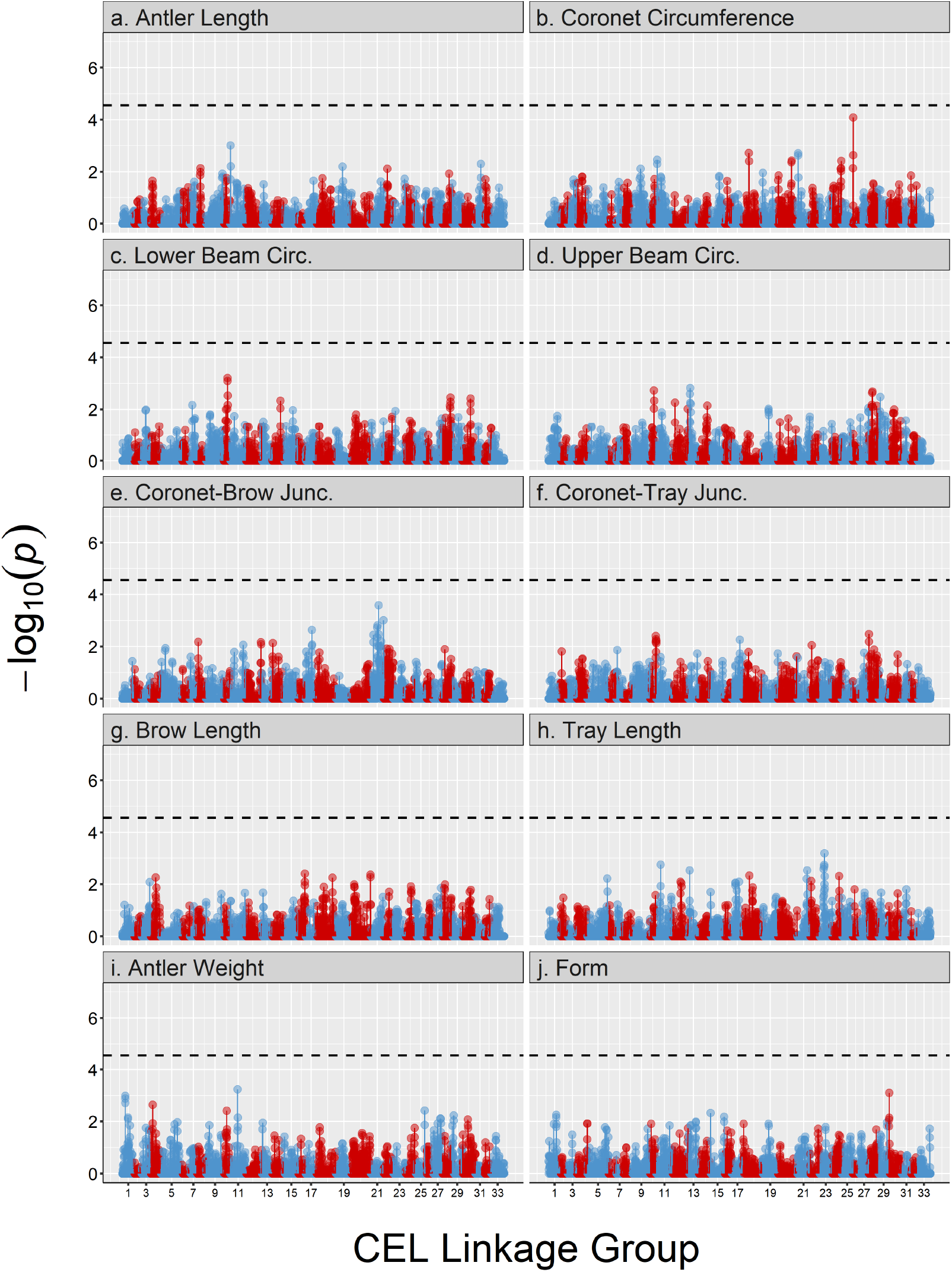
Regional heritability analysis for antler measures. The dashed line indicates the significance threshold equivalent to *α* = 0.05. Points are colour-coded by chromosome. Underlying data is provided in Table S6

### Distribution and quantification of SNP effect sizes

A total of 897 unique SNPs had significant non-zero effects across the ten antler measures (Table 4; full results are provided in Table S11). The number of significant non-zero SNPs ranged from 15 SNPs (antler length) to 279 SNPs (lower beam circumference; Table 4), although antler weight and upper beam circumference had no significant non-zero effect SNPs. Several SNPs showed pleiotropic effects i.e. they were associated with more than one measure (Table S11), with the underlying proportion ranging from 6% (coronet-tray junction) to 68% (tray length). Lower beam circumference and antler form showed distributions that included more extreme SNPs with large effects on phenotype outside the 5% to 95% quantile boundaries, whereas most other measures had more uniformly distributed effect sizes (Table 4). For the 11 antler PCs, only 159 unique SNPs had non-zero effects (Table S10; full results in Table S12). No pleiotropic SNPs were observed, most likely due to the independence of each PC. PC4 and PC9 had no non-zero effect SNP associations. Only about 25% of the 159 SNPs were in common with the 897 SNPs in the antler measure analysis. This was mainly due to the higher order PCs (PC6 to PC11) sharing no SNP associations with any antler measures, while other PCs had similar or more numbers of shared and unique SNPs (PC1 and PC2).

**Table 4:**
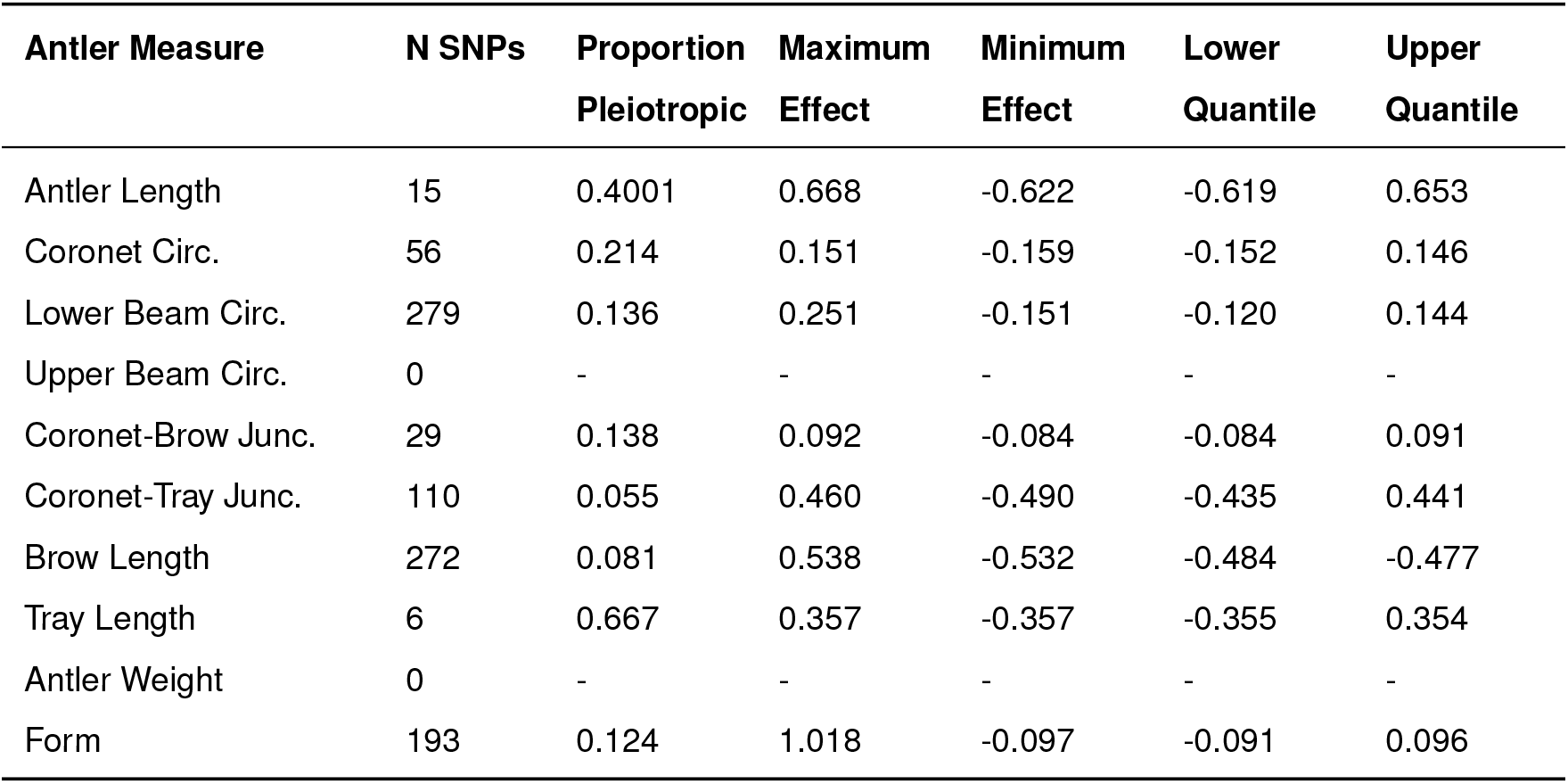
Summary of SNPs with non-zero effects on antler measures. N SNPs is the number of SNPs with non-zero effects. Proportion pleiotropic is the proportion of SNPs with non-zero effects on other antler measures. Maximum and minimum effects are given relative to the data scale (units = cm for lengths, g for weight). Lower and upper quantiles refer to the 5% and 95% boundaries of the effect size distribution respectively.

Some of the non-zero effect SNPs with large effect size estimates after FDR explained large proportions of overall genetic and phenotypic variance for their respective traits. The SNP *cela1_red_2_101997097* explained about 17% of the overall additive genetic and 4% of phenotypic variance in antler length (SNP variance = 2.609, SE = 1.094), while for PC8 (which is strongly representative of antler length; Figure 2) the marker *cela1_red_4_41756873* explained 39% of the additive genetic and 4% of the phenotypic variance (SNP variance = 0.019, SE = 0.008). Despite these seemingly large effect marker associations the standard errors for the SNP variance estimates were generally large (see Tables S13 and S14 for summaries of variances, and Tables S15 and S16 for full results for the antler measures and PCs, respectively). Overall, as the uncertainty around the importance of the SNP effects is large and none of these marker achieved genome-wide significance in any of the GWAS, the influence of specific SNP loci on variation in antler measures and PCs must be interpreted with caution.

## Discussion

In this study, we have used a genomic approach to determine the genetic architecture of antler morphology in the red deer of Rum. We have shown that antler morphology is heritable and repeatable over an individual’s lifetime, and that it is likely to be highly polygenic with a moderate degree of shared genetic architecture across different antler traits. Genome-wide association and regional heritability studies failed to identify any genomic regions associated with antler measures and their principal components, with the exception of a single region on linkage group 21 associated with variation in PC9. Most traits were underpinned by SNPs with uniformly small effect sizes, whereas others, such as coronet circumference and antler form, showed distributions that included some SNPs with larger effects on phenotype. Our findings suggest that antler morphology has a highly polygenic architecture with many loci of small effect. Here, we discuss how our findings build on previous quantitative genetic studies in the Rum red deer system, and how they inform the broader question of the distribution of genetic architectures of sexually selected male weaponry and the consequences for its evolution.

### Heritability and repeatability of antler morphology

All 10 antler measures were significantly heritable (ranging from 0.229 to 0.414, Table 2) and were similar to previous heritability estimates for antler weight and form in the same population (Kruuk *et al.*, 2014). The strong agreement of estimates from both the pedigree and GRM approaches indicate that the SNPs present on the red deer SNP chip are in sufficiently high LD with causative loci to allow accurate estimation of trait heritabilities in this population (Yang *et al.* 2010; Figure 3). Indeed, LD is maintained at a relatively constant level over a distance of up to 1Mb, after which it starts to slowly decay (Figure S1). All antler traits were also highly repeatable, with between 38.5% and 66.1% of the phenotypic variance explained by additive genetic, permanent environment and birth year effects. This indicates that antler morphology is temporally stable over an individual’s adult life, despite the annual shedding and regrowth of antlers. Similar findings were found for the 11 principal components, with lower order PCs generally exhibiting higher heritabilities and repeatabilities (Table S2); we discuss why we think this is the case in the section on genetic correlations and constraints below. It should be noted that the heritabilities may be slightly over-estimated due to any common environments on a fine spatio-temporal scale that cannot captured by the animal models (Stopher *et al.*, 2012). These findings support previous work showing that traits under sexual selection can have substantial underlying genetic variation in wild populations (Kruuk *et al.*, 2008; Merila *et al.*, 2001; Pomiankowski & Moller, 1995), where morphological traits often show heritabilities of a similar magnitude even when associated with fitness (Johnston *et al.*, 2013; Bérénos *et al.*, 2014; Bourret *et al.*, 2017; Malenfant *et al.*, 2018).

### The polygenic architecture of antler morphology

Genome-wide association studies and regional heritability analyses across all antler traits and PCs showed no significant associations, with the exception PC9 (discussed in the next section). Generally, GWAS can only detect loci with moderate to large effects on phenotype and which only partially explain the trait heritabilities, with the remainder termed the “missing heritability” (Manolio *et al.*, 2009; Golan *et al.*, 2014). For example, a meta-analysis of human GWAS found that the heritability attributed to all common SNP variants was significantly higher than that of the SNPs that achieved genome-wide significance. (Shi *et al.*, 2016), meaning that large numbers of “non-significant” SNPs will contribute to the additive genetic variation. To characterise the genetic architecture beyond GWAS alone, we employed two additional approaches. The first, regional heritability (Nagamine *et al.*, 2012), incorporated the haplotypic diversity within genomic regions. This approach detected a large contribution of defined genomic regions to a single PC, but not to any of the other antler measures or PCs. The second, the Empirical Bayes false discovery rate and effect size estimation, incorporated information from the GWAS effect size estimates and their error to show that a substantial number of SNPs had non-zero effects on antler morphology (Stephens, 2016). Taken together, these analyses provide compelling evidence that most aspects of antler morphology have a highly polygenic architecture (Fisher, 1930; Barton *et al.*, 2017). This in line with findings from other studies of wild organisms that have identified polygenic architectures for morphological and life history traits (Robinson *et al.*, 2013; Santure *et al.*, 2013; Pallares *et al.*, 2014; Berenos *et al.*, 2015; Husby *et al.*, 2015). In this study, we modelled antler morphology using both independent measures of antler morphology and a principal component framework to characterise different dimensions of shape variation. An advantage of using both approaches was that it allowed us to identify a greater number of potentially causal loci, which supports the usefulness of this approach when trying to characterise the genetic architecture of a complex morphological trait (such as in Pallares *et al.* 2014).

### Genomic regions associated with antler morphology

Three genomic windows within the CEL linkage group were associated with PC9 at the genome-wide level. The main contributing antler measure to PC9 is coronet circumference The region covered by the most highly associated window explained ~66% of the additive genetic variance, representing a large part of the overall heritability estimated using the whole GRM (~19%). The region contains a number of candidate genes among which are gasdermin C (*GSDMC*); MYC proto-oncogene (*MYC*); ArfGAP with SH3 domain, ankyrin repeat and PH domain 1 (*ASAP1)* and cellular communication network factor 4 (*CCN4*). Whilst none have previously been implicated in antler morphology, they have associated functions that make them potential candidate genes. Both the enhancer protein *ASAP1* and *CCN4,* which is a type of connective tissue growth factor, are linked to bone ossification and bone cell differentiation in mice (The Jackson Laboratory, 2019; Schreiber *et al.*, 2019; Maeda *et al.*, 2015), processes which are likely to be vital to antler regeneration, rapid growth and the ability of antlers to withstand impact (Goss, 1983). Upregulation of *GSDMC* is implicated in carcinogenesis in mice, as a consequence of an interrupted growth factor signalling pathway (Miguchi *et al.*, 2016) and *MYC* is a potent oncogene that is implicated in many human cancers and promotes rapid tumor cell proliferation (Beroukhim *et al.*, 2010; Lin *et al.*, 2012); recent work suggests that rapid regeneration of bony antlers has evolved by upregulating cell proliferation pathways while suppressing tumorigenesis (Wang *et al.*, 2019).

Despite being linked to compelling candidate gene regions, with our current data it is virtually impossible to determine exactly which genes in this region drive the association with PC9. Furthermore, as PC9 only represents ~4% of the overall phenotypic variance among all antler PCs, we expect effect sizes of causal loci to be very small. Validating the findings for this association would be challenging, as replication of a similar PC in other deer populations would have to consider its main contributing antler measures (e.g. coronet circumference); additionally, the same variants may not be associated with this trait in different populations. A more feasible approach may be to type a higher density of SNP loci to characterise more variation at (or in tight linkage) with potential causal loci in the Rum deer population.

### Genetic correlations and constraints on antler morphology

Almost all antler traits were positively genetically correlated, with the exception of the coronet-tray junction, which was negatively genetically correlated with tray length, lower beam circumference and antler form. These findings were reflected by the PC analysis, where the PC explaining the most variance in antler morphology (PC1, ~41%) combined equal information from all antler measures, whereas those explaining declining amounts of variation represented one or two antler traits. The Empirical Bayes analysis identified varying degrees of marker pleiotropy associated with the antler measures (Table 4) that were consistent with the observed genetic correlations. Nevertheless, there were some exceptions, such as for brow length and antler weight, which both showed strong positive genetic correlations yet had small proportions of shared loci (brow length) or no associated loci at all (antler length). This incongruity may be explained by the large variation in the number of non-zero effect SNPs detected between antler measures, which could be due to differences in the effect size distribution of markers.

The large heritabilities of individual antler traits suggest there is potential for response to selection, but our results further add to previous findings that constraints at the genetic level may affect how the population may respond to selection. Previous work showed that a large part of genetic variance in antler weight is not available to selection due to the lack of genetic covariance between weight and fitness (Kruuk *et al.*, 2002, 2014). In the current study, antler weight was highly correlated with most other antler measures on a genetic level, which may likely indicate that large parts of the genetic variance of these antler measures are also unavailable for selection, thus limiting the evolutionary potential of antler morphology despite large heritability estimates. In the PC analysis, higher order PCs explained much smaller amounts of variation and had very low additive genetic variation. It is possible that these PCs could represent morphologically stable aspects of the antlers, where lower heritabilities could indicate past strong stabilising selection on the combination of trait aspects represented by that PC. Other studies exploring multivariate sexual selection on a suite of traits found that despite large genetic variance in univariate analyses, there can be very little genetic variance available in the trait composition that is the target of selection (Hunt *et al.*, 2007; Van Homrigh *et al.*, 2007).

The discovery that the genetic architecture of antler morphology is highly polygenic also suggests that further evolutionary mechanisms may be partially responsible for the maintenance of genetic variation in this trait. As discussed in the introduction, traits with many genes of small effect can present a large mutational target for the introduction of novel genetic variation which can contribute to the genetic variation in a trait (Rowe & Houle, 1996). Consequently, although selection on polygenic traits can lead to rapid changes in trait mean, under the infinitessimal model the distribution of underlying genetic effects is expected to remain relatively constant, counteracting the loss of genetic variation. (Barton *et al.*, 2017; Sella & Barton, 2019). Finally, pleiotropic effects of loci that share a similar complex architecture with other traits and/or are in LD with loci associated with fitness could maintain genetic variation through conflicts and trade-offs (Lande, 1982).

## Conclusions

In this study, we have shown that antler morphology is heritable, has a polygenic genetic architecture, and some degree of shared genetic architecture between different antler measures. A single region association is linked to candidate genes that could potentially have an effect on antler morphology, but more work would be required to validate this finding in this and other populations. Future work in this system will integrate knowledge of the genomic architecture of antler morphology with fitness measures to further dissect constraints on trait evolution within this population. Ultimately, our findings corroborate the expectation for a quantitative trait such as multidimensional weaponry traits to conform to a polygenic genetic architecture of many genes with small effects.

## Author Contributions

L.P. and S.E.J. designed the study. J.M.P and L.K provided the data. L.P., J.H and S.E.J. analysed the data. L.P. and S.E.J. wrote the first draft of the paper and all authors contributed to revisions.

## Acknowledgements

We thank Alison Morris, Sean Morris, Martin Baker, Fiona Guinness, Tim Clutton-Brock and many others for collecting field data and DNA samples over the course of the long-term study. We thank Scottish Natural Heritage for permission to work on the Isle of Rum National Nature Reserve. Andy Arthur kindly provided Figure 1. Philip Ellis prepared samples for DNA extraction and the Wellcome Trust Clinical Research Facility Genetics Core in Edinburgh performed the genotyping. This work made extensive use of the University of Edinburgh Compute and Data Facility (http://www.ecdf.ed.ac.uk/). The long-term project on Rum red deer is funded by the UK Natural Environment Research Council (NERC) and the SNP genotyping was funded by a European Research Council Advanced Grant to J.M.P.. L.P is supported by a NERC E3 Doctoral Training Programme PhD Studentship. S.E.J. is supported by a Royal Society University Research Fellowship.

