## Supplementary figures and images for "Genomic analysis reveals a polygenic architecture of antler morphology in wild red deer (*Cervus elaphus*)"

### Supplementary figure 1

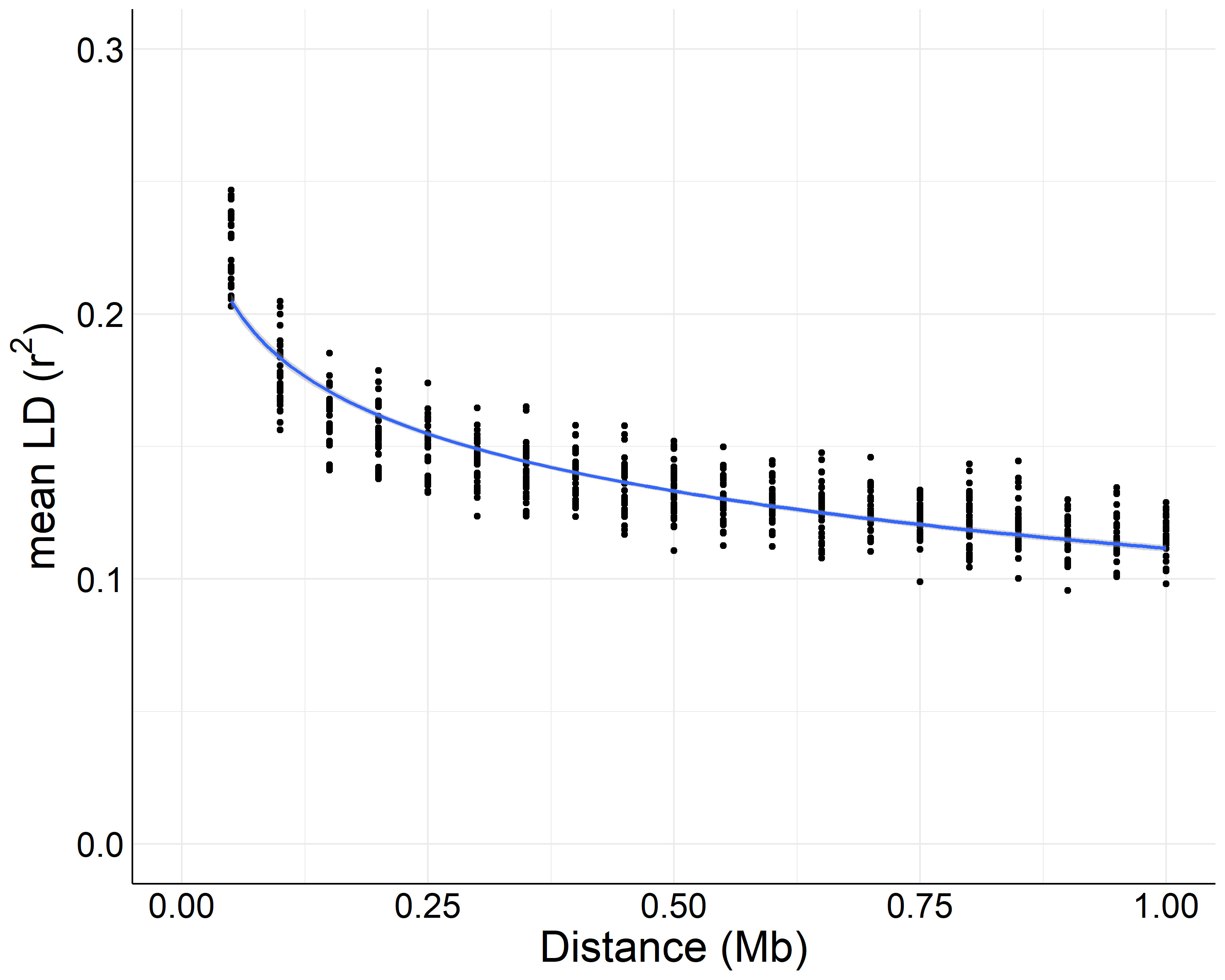

### Supplementary Figure 2

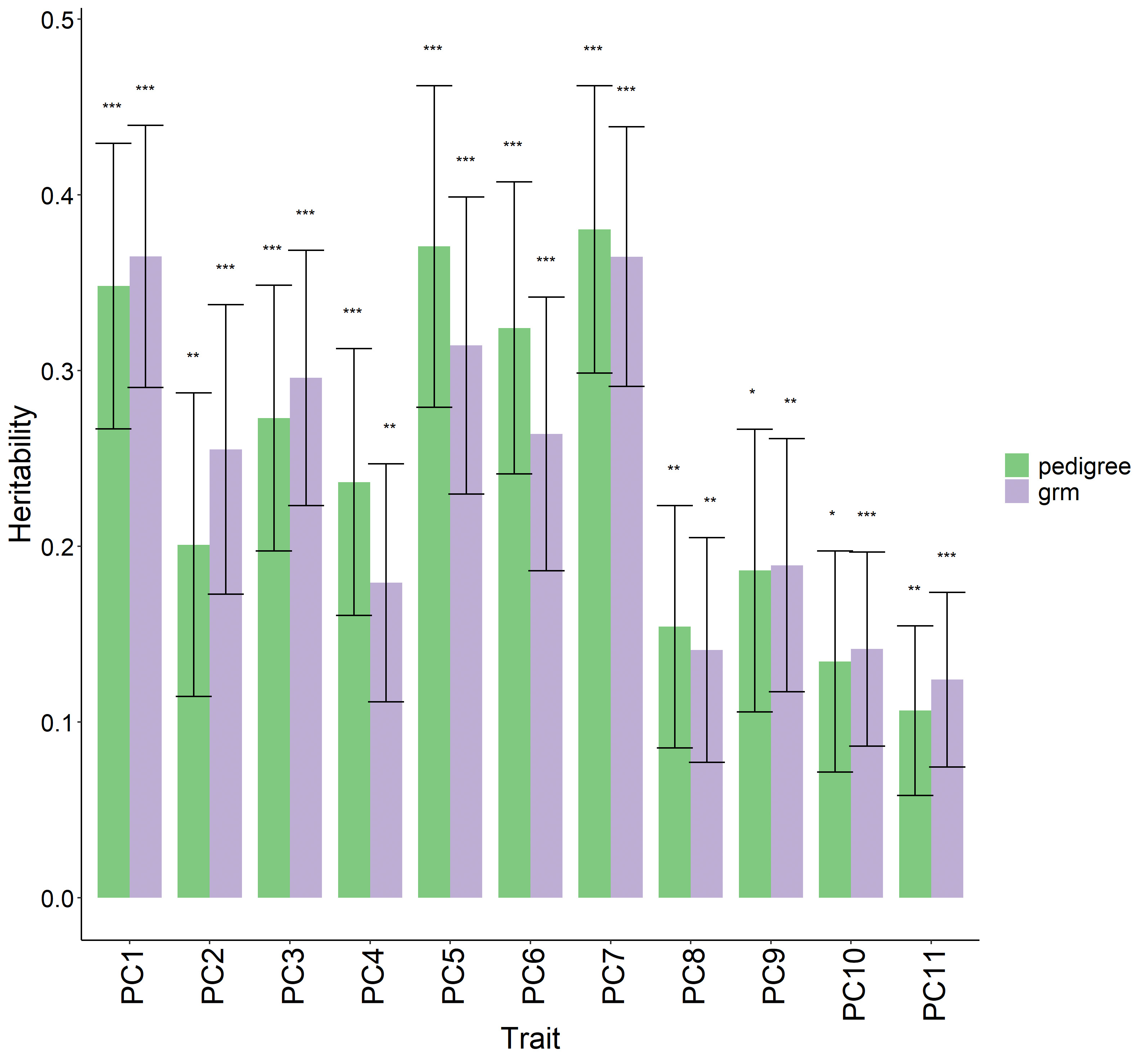

### Supplementary Figure 3

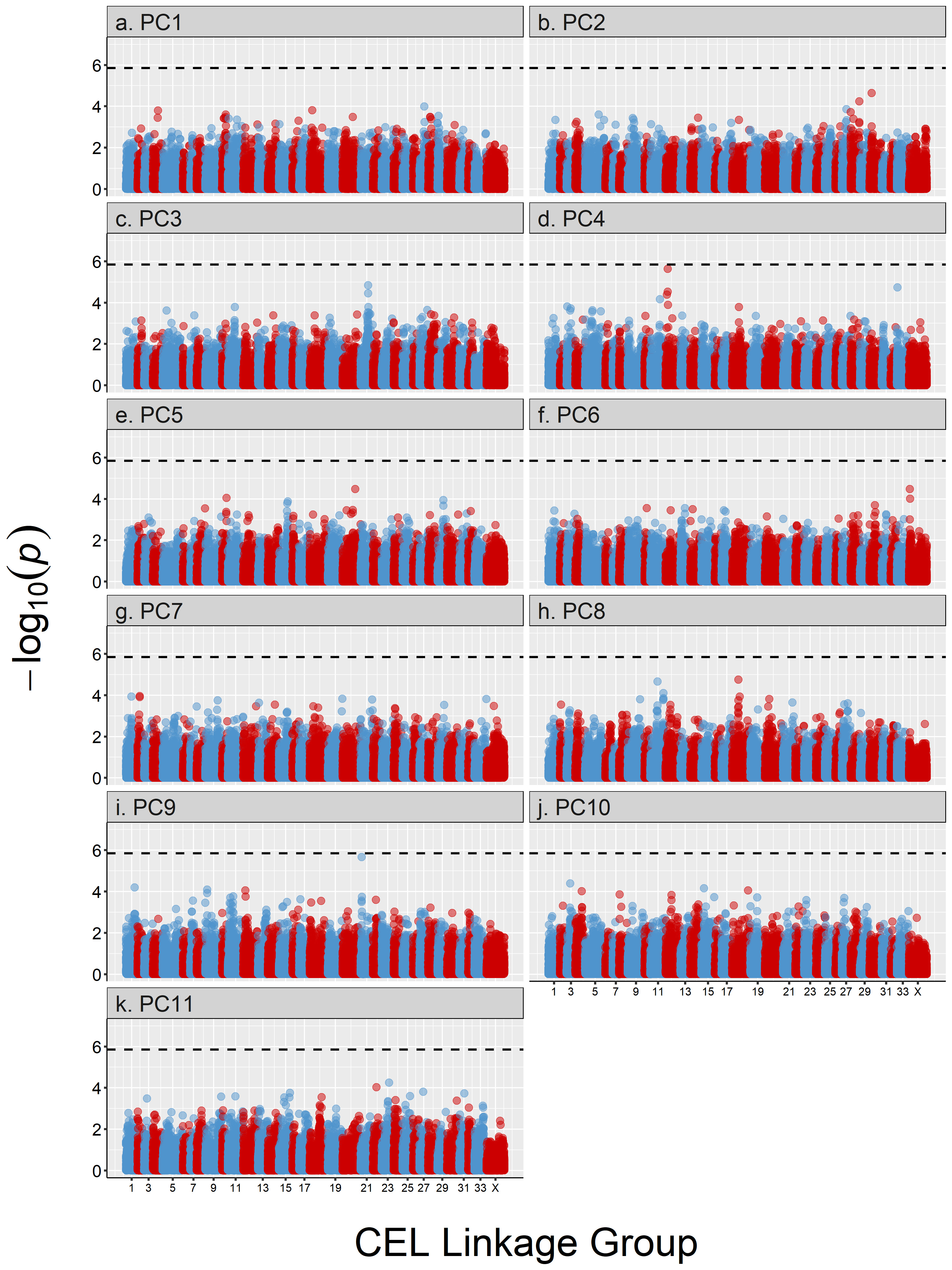

### Supplementary Figure 4

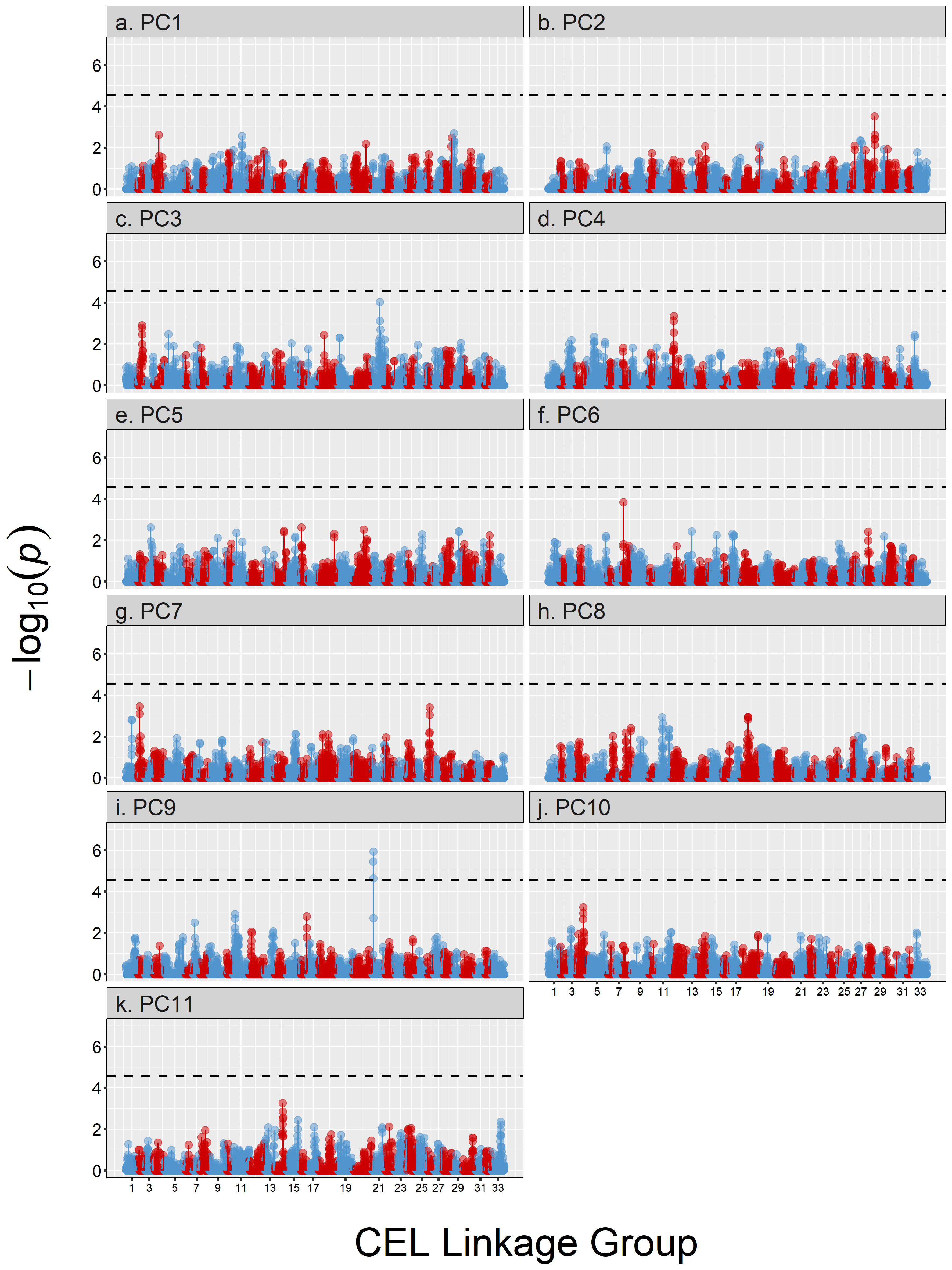
